# Targeted modification of gene function exploiting homology-directed repair of TALEN-mediated double strand breaks in barley

**DOI:** 10.1101/019893

**Authors:** Nagaveni Budhagatapalli, Twan Rutten, Maia Gurushidze, Jochen Kumlehn, Goetz Hensel

## Abstract

Transcription activator-like effector nucleases (TALENs) open up new opportunities for targeted mutagenesis in eukaryotic genomes. Similar to zinc-finger nucleases, sequence-specific DNA-binding domains can be fused with effector domains like the nucleolytically active part of FokI in order to induce double strand breaks (DSBs) and thereby modify the host genome on a predefined target site via non-homologous end joining. More sophisticated applications of programmable endonucleases involve the use of a DNA repair template facilitating homology-directed repair (HDR) so as to create predefined rather than random DNA sequence modifications. The aim of this study was to demonstrate the feasibility of editing the barley genome by precisely modifying a defined target DNA sequence resulting in a predicted alteration of gene function. We used *gfp*-specific TALENs along with a repair template that, via HDR, facilitates conversion of *gfp* into *yfp* which is associated with a single amino acid exchange in the gene product. As a result of co-bombardment of leaf epidermis, we detected YFP accumulation in about 3 out of 100 mutated cells. The creation of a functional *yfp* gene via HDR was unambiguously confirmed by sequencing of the respective genomic site. Predictable genetic modifications comprising only a few genomic base pairs rather than entire genes are of particular practical relevance, because they might not fall under the European regulation of genetically engineered organisms. In addition to the allele conversion accomplished *in planta,* a readily screenable marker system is introduced that might be useful for optimization approaches in the field of genome editing.

## INTRODUCTION

Transcription activator-like effector nucleases (TALENs) offer exciting opportunities for targeted mutagenesis. Like the zinc-finger nucleases (ZFNs), the TALENs comprise a target sequence-specific DNA-binding domain fused with the nucleolytically active part of endonuclease FokI that induces the formation of double strand breaks (DSBs); random mutations arise at the target site as a result of error-prone DNA repair effected via non-homologous end joining (NHEJ). First established in yeast and human cell lines, numerous other genomes are now known to be amenable to this mode of genome modification (Joung and Sander 2013). ZFNs, TALENs and the more recently developed RNA-guided endonuclease (RGEN, also called CRISPR/Cas9) system (for review see Voytas 2013) have yet to be widely exploited in plant genomes, but it has recently been demonstrated that heritable TALEN-mediated mutagenesis can be achieved with a relatively high efficiency in barley (Gurushidze *et al*. 2014). An attempt to use TALEN and RGEN technology to mutagenize the orthologs of the barley susceptibility factor MLO in hexaploid wheat resulted in around 6% of plants mutated in at least one of the three homoeologous genes (Wang *et al*. 2014). An advantageous feature of the TALENs is the high modularity of the nucleotide-binding repeats, which allows for the assembly of binding domains targeting almost any site of choice in the host genome (Reyon *et al*. 2012; Schmidt-Burgk *et al.* 2013). Since FokI activity is only manifested as a dimer, two TALEN units need to be expressed simultaneously. Consequently, the target site comprises about 50 nt, sufficient to ensure a high degree of specificity.

The inclusion of a DNA repair template facilitates homology-directed repair (HDR), which has been shown to increase the chances of effecting predefined rather than random DNA sequence modifications. The NHEJ repair frequency of DSBs induced within a reporter gene, as generated by ectopically expressing I-*SceI* endonuclease in barley, lay in the range 47-58%, including some 1-3 nt indels (Vu *et al*. 2014). A further 1-8% of the modifications displayed synthesis-dependent strand annealing associated with NHEJ, while 20-33% arose from homologous recombination. The surprisingly high proportion of HDR has encouraged the possibility of deploying a repair template-mediated genome editing approach.

In plants, homologous recombination-based modifications have been largely limited to alterations resulting in a selectable or a screenable phenotype (reviewed by Voytas 2013). Recent examples of sequence knock-in (targeted insertion) employing RGEN-mediated DSB induction have been reported in both rice and *Nicotiana benthamiana* protoplasts (Li *et al*. 2013; Shan *et al*. 2013), in which the templates were between 72 bp and 647 bp in length; the resulting rate of incorporation of restriction enzyme recognition sites into the target loci was 6-10%. More recently, Schiml and colleagues (2014) achieved heritable genome modifications by integrating a resistance gene cassette into an endogenous *ADH1* locus via *in planta* gene targeting (GT) after RGEN-mediated DSB induction. In this context, these authors suggested that the directed exchange of single base pairs within the coding sequence of a protein of interest is one of the challenges still needed to be addressed using *in planta* GT technology. Baltes *et al*. (2014), working with *Nicotiana tabacum*, used circular geminivirus replicons (GVRs) to introduce sequence-specific nucleases as well as DNA repair template, and showed that ZFN, TALEN and RGEN elements could all be expressed and then induce mutation at a high frequency. A non-functional *GUS:NPTII* transgene was corrected by transforming tobacco leaves with these GVRs reconstituting a functional GUS copy. However, so far, repair template-mediated conversion of a functional allele towards the gain of altered gene functionality has not yet been reported for any plant host.

Here we exploit TALEN technology involving HDR of DSBs to generate a precisely directed genome modification at the individual nucleotide level *in planta*. This approach represents a novelty in monocotyledons which are of utmost agricultural importance. To exemplify the principle, a stably integrated *gfp* gene is converted into *yfp*, which is readily screenable at the cellular level. The two gene products differ only in a single T203Y amino acid which causes an altered light emission spectrum (Wachter *et al*. 1998). We show that yellow fluorescing cells can be reproducibly generated via allele conversion and unambiguously confirm the genetic modification by sequencing the genomic target site.

## MATERIALS AND METHODS

### Plant materials

The two transgenic winter barley (*Hordeum vulgare*) cv. ‘Igri’ lines, denoted BPI 09 and BPI 11, each carry a single copy of *gfp* (Fig. 1A). The grain was germinated (14/12°C day/night, 16 h photoperiod at 20,000 lux), and the seedlings vernalized for eight weeks (4°C, 9 h photoperiod), then raised in a glasshouse (18/14°C, 16 h photoperiod at ∼25,000 lux). Spikes were harvested when the awns had just emerged from the flag leaf, as described by Kumlehn *et al*. (2006).

### Plasmids and transgenic barley plants

A pair of *gfp*-specific TALEN constructs pGH297 (carrying the left-hand unit) and pGH400 (right-hand unit) was generated as described by Gurushidze *et al*. (2014) (Supporting Information, Figure S1A, B). A fragment comprising *yfp* gene along with *NOPALINE SYNTHASE* terminator were synthesized and introduced into EcoRV cloning site in pUC57 at GenScript (Piscataway, NJ, USA), then subsequently cloned at *Bst*XI/*Bam*HI sites into pYF133 (Fang *et al*. 2002) (pGFP in this study, Figure S1C) to generate pGH418, in which *gfp* was replaced by *yfp* (pYFP, in this study, Figure S1D). To avoid the possibility of the *gfp*-TALEN induced mutation of *yfp*, its binding sites were modified by introducing synonymous codons, leaving the polypeptide sequence unaltered (Figure 1B). Electroporation was used to introduce the TALEN constructs into *Agrobacterium tumefaciens* strain LBA4404/pSB1, which harbors the disarmed Ti plasmid pAL4404 and the hypervirulence-conferring plasmid pSB1 (Komari *et al*. 1996). Microspore isolation and the agroinoculation of embryogenic BPI 09 and BPI 11 pollen with each of the two *gfp*-specific TALEN constructs were carried out following Kumlehn *et al*. (2006), except that the population density of the transformed LBA4404/pSB1 cells was adjusted to 2.5 × 10^7^ colony-forming units per mL. Bialaphos was used as the selective agent. The regenerants were screened for the presence and expression of FokI nuclease and two lines showing the expression of the right- (335R and 338R) and another two lines of the left-hand (319L and 462L) *gfp*-TALEN units were carried forward.

**Figure 1.**
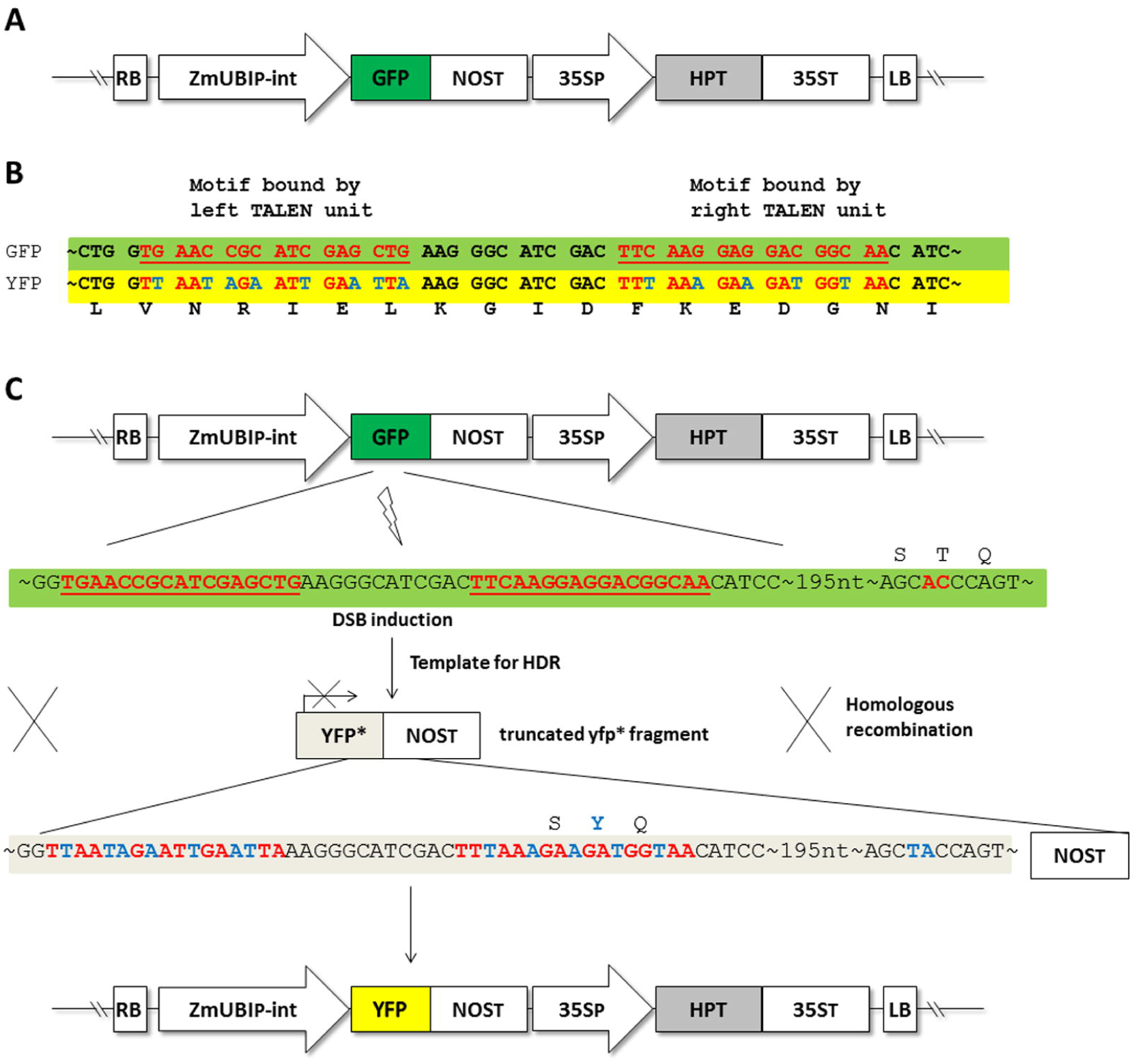
Concept of HDR of TALEN-induced DSBs in barley. (A) Barley genomic region with T-DNA of *gfp* plants used for re-transformation with *gfp*-specific TALEN constructs. (B) *gfp*-specific TALEN target site (with DNA-motifs bound by the left and right TALEN units indicated in red and underlined) aligned with the respective sequence of the truncated *yfp* fragment (*yfp*\*) used as a template for HDR, which is modified by substitution with synonymous codons (in blue) to prevent the TALENs from binding, while the YFP amino acid sequence is retained. (C) Mechanism showing the *gfp* TALEN-induced DSBs in T-DNA region of *gfp* and used truncated *yfp*\* fragment as template for HDR, leading to conversion of *gfp* to *yfp*.

### Biolistic transformation of leaf cells

Gold particles of diameter 1.0 μm (Bio-Rad, Munich, Germany) were coated with DNA, and a 1.5 mg aliquot used for each shot. The concentration of coating DNA was 1 μg/μL (pGH297, pGH400) and 2-3 μg/μL (834 bp *Btg*ZI/*Bss*HII-linearized and gel-purified pUC57-*yfp* DNA fragment), precipitated from solution according to Sanford *et al*. (1991). The transformation was effected using a Biolistic Particle Delivery System (PDS)-1,000/He device equipped with 1,100 psi rupture disks (Bio-Rad, Munich, Germany). Five primary leaves sampled from 7-10 day old seedlings were laid adaxial side up on 1% agar containing 20 μg/mL benzimidazol and 20 μg/mL chloramphenicol. After bombardment, the leaf material was held at room temperature for 17-20 h before being subjected to fluorescence microscopy.

### Biolistic transformation of embryo cells

Immature grains (12-16 days after anthesis) were surface sterilized by immersion for 30 min in 70% (v/v) ethanol, followed by 20 min in 2.4% (w/v) sodium hypochlorite plus 0.1% (v/v) Tween 20 and four rinses in sterile distilled water. The embryos were dissected out aseptically and cultured for five days at 24°C in the dark on a solid (3 g/L phytagel) medium containing 4.4 g/L Murashige and Skoog salts, 112 g/L B5 vitamins, 30 g/L sucrose, 5 µM copper sulphate and 5 mg/L dicamba (all chemicals were purchased from Duchefa, Haarlem, The Netherlands). Prior to bombardment, the callus was transferred onto the same medium supplemented with 0.4 M sorbitol for 4 h. A 3 mg aliquot of DNA-coated gold particles was used to bombard 20 calli, employing the same device as above, but using a 900 rather than a 1,100 psi rupture disk. After bombardment, the material was held for 16-18 h at 24°C in the dark before being subjected to fluorescence microscopy.

### Confocal microscopy

The DSB induction efficiency caused by the TALEN constructs and that of their subsequent HDR was determined by establishing the ratio of *yfp*-expressing and *gfp*-non-expressing cells in five 2×1 cm leaf segments. Only cells from the stomatal cell rows (guard cells and adjacent epidermal cells, see Figure S2) were counted. GFP (488 nm excitation, 491-530 nm emission) and YFP (514 nm excitation, 517-560 nm emission) signals were captured by confocal microscopy, and the identity of the fluorophores was confirmed by photospectrometric analysis.

### Genomic DNA isolation and PCR

Genomic DNA was isolated from snap-frozen transiently transformed leaves following the Palotta *et al*. (2001) protocol. Each subsequent 20 µL PCR contained 50-100 ng template, and was based on the primer pairs listed in Table S1 and standard PCR conditions. The reaction products were purified using a QIAquick PCR Purification kit (Qiagen, Hilden, Germany) for subsequent sequencing.

### RNA isolation and reverse transcription

Total RNA was isolated using the TRIzol reagent (Life Technologies, Darmstadt, Germany), following the manufacturer’s protocol. After DNase I treatment (DNA-free kit, Life Technologies, Darmstadt, Germany), the first cDNA strand was synthesized using an iScript Select cDNA synthesis kit (Bio-Rad, Munich, Germany). The resulting cDNA served as the template for subsequent RT-PCRs based on the FokI domain primer pair (Figure S1E).

## RESULTS AND DISCUSSION

Our aim here was to demonstrate the feasibility of editing a plant genome by precisely modifying a defined target DNA motif. A *gfp*-specific TALEN was used in conjunction with a repair template which, via HDR, was intended to convert *gfp* into *yfp*, an alteration achievable by inducing a T203Y substitution (Figure 1C). Such an alteration would be readily detectable, as it produces a shift in fluorescence emission from 509 nm (GFP) to 527 nm (YFP) (Wachter *et al*. 1998). In an initial experiment, immature embryos of barley transgenic lines harboring *gfp* (Figure 1A) and actively expressing either (1) the left-hand *gfp*-TALEN unit (line 462L) or (2) the right-hand *gfp*-TALEN unit (line 335R) (Figure S1E) were co-bombarded with vectors carrying the complementary *gfp*-TALEN unit (pGH400 and pGH297, respectively, Figure S1A, B) and a linearized, truncated, non-functional *yfp*\* fragment (Figure 1C). The design of the *gfp*-TALEN pair and the generation of transgenic plants essentially followed Gurushidze *et al*. (2014). In order to prevent TALEN-mediated mutations in the repair template, its TALEN recognition sequences were modified by substitution with synonymous codons, which retained the YFP amino acid sequence (Figure 1B). One day after bombardment a number of yellow fluorescing cells were detected (Figure S3A, E) and confirmed by corresponding emission peaks recorded by the confocal laser scanning microscope (Figure S3B). Some of the cells lack GFP fluorescence while showing YFP activity (Figure 3C-E).

To assess the frequency of allele conversion, leaves of the homozygous *gfp* barley plants additionally expressing either the left-hand (lines 319L and 462L) or the right-hand (lines 335R and 338R) members of the *gfp*-TALEN pair were bombarded as above. After bombardment, 2 cm long leaf segments were analysed with regards to the proportion of the total number of cells (stomatal complex and the first layer of adjacent epidermal cells), the non-fluorescent cells and the ones showing yellow fluorescence (Figure S2). Possible scenarios in treated cells with regards to their fluorescence are detailed in Table 1.

**Table 1.** Possible scenario after bombardment of leaf material

| Scenario | Fluorescence |
| --- | --- |
| (G1) non-transformed cells (not met by a particle) | Green |
| (G2) transformed cells where TALENs did not effect a DSB in the target site |  |
| (G3) cells with **both *gfp* alleles** being **unaltered owing to correct repair** or to a mutation not affecting gene function |  |
| (G4) cells with **one *gfp* allele** being functionally affected **via NHEJ, whereas the other one** is unaltered or its function is retained despite of a (minor) mutation |  |
| (GY1) cells with **one *gfp* allele** being modified by **HDR using the YFP-repair template, whereas the other one** remained functionally unaffected | green and yellow |
| (Y1) cells with both *gfp* alleles being modified by **HDR using the YFP-repair template** | yellow |
| (Y2) cells with **one *gfp* allele** being modified by **HDR using the YFP-repair template, whereas the other one** is functionally affected by NHEJ |  |
| (N1) cells with both *gfp* alleles being functionally affected by NHEJ | no |
| (N2) cells that have not survived |  |

After bombardment of negative control samples using the very same *gfp*-TALEN unit as was already stably integrated in the plant material used (e.g. bombardment of line 338R using pGH400 also carrying the right TALEN unit), ca. 1% (207 out of 20,238 cells) of the cells failed to show green fluorescence, suggesting these to represent the rate of cell death due to mechanical damage (Figure 2E, dark grey bars). A further ca. 2% (495 out of 19,141 cells) could be accounted for by random mutagenesis at the TALEN target site when complementary *gfp*-TALEN units were used without the *yfp*\* repair template (Figure 2E, light grey bars) (e.g. bombardment of line 319L with pGH400 carrying the right TALEN unit). YFP fluorescence was only observed when the treatment involved the truncated *yfp*\* repair template and both complementary *gfp*-TALEN units (Table S2; Figure 2C). Under this condition, on average 2-3% of the cells mutated at the target site were found to express *yfp*, thereby displaying a conversion of at least one *gfp* allele into *yfp*, which is a reasonable success rate of precise genome editing (Table S2).

**Figure 2.**
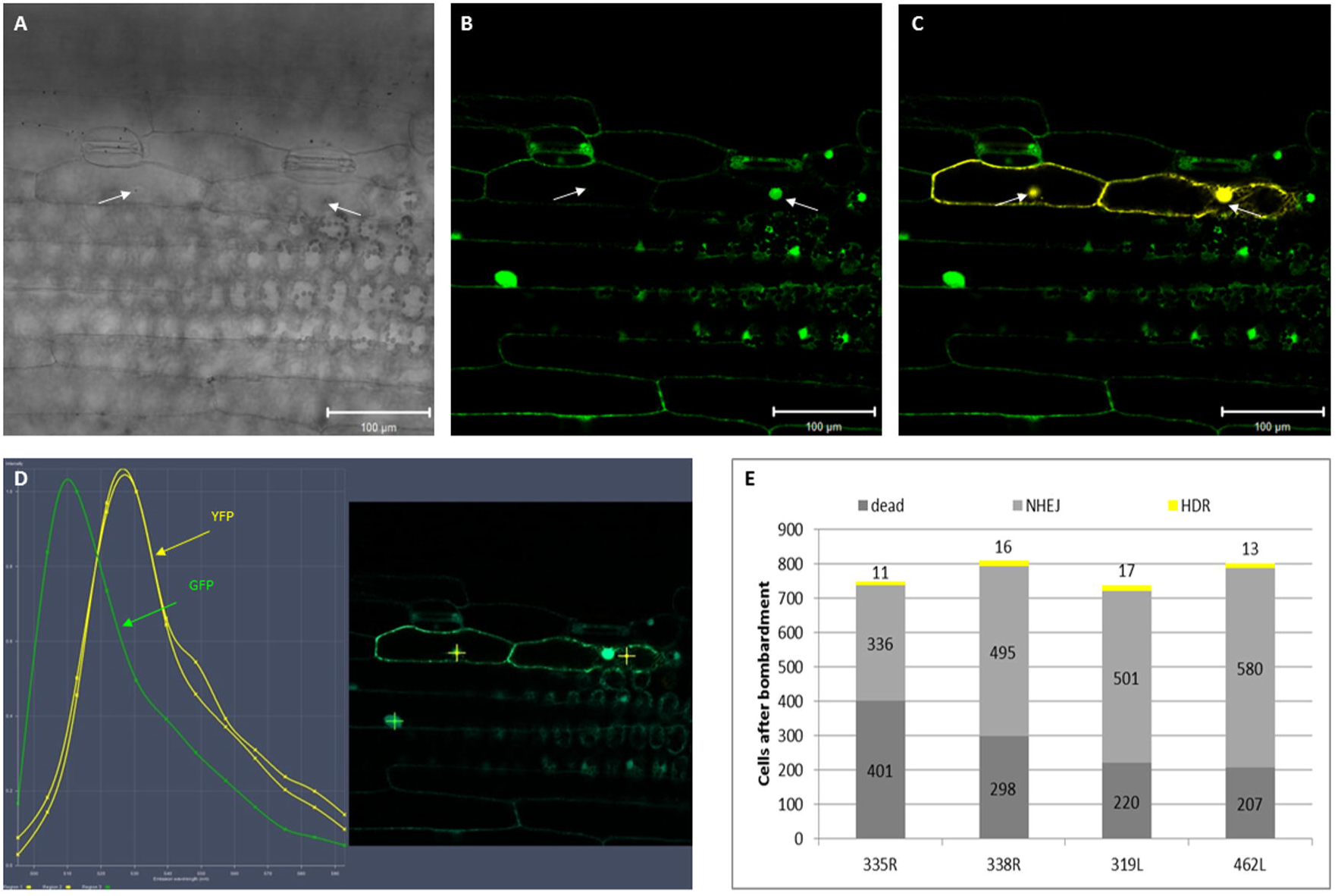
HDR following the induction of TALEN-mediated DSBs in barley. (A) Bright field of a leaf of line 319L (carrying *gfp* and the left-hand TALEN unit) taken 20 h after bombardment with the right-hand TALEN unit and linearized *yfp*\* fragment. (B) Epifluorescence of the same leaf shown in (A) following excitation with 488 nm laser light and the spectral unmixing of GFP using confocal microscopy. (C) Epifluorescence of the same leaf shown in (A) following excitation with 488 nm laser light and the spectral unmixing of GFP (green) and YFP (yellow) signals using confocal microscopy. Bright field, GFP and merged GFP and YFP signals. Arrows indicate nuclei of two adjacent epidermal cells. Bar: 100 μm and (D) Lambda stack of same materials shown in (A-C) used to visualize the presence of GFP (emission peak at 509 nm) and YFP (527 nm). (E) The bombardment of lines 319L and 462L (carrying *gfp* and the left-hand TALEN unit), and 335R and 338R (gfp plus the right-hand TALEN unit). Bars represent the number of non-fluorescing cells resulting either from mechanical damage (dark grey, numbers inferred from control bombardments in the absence of the complementary TALEN unit) or from TALEN-mediated DSB followed by error-prone NHEJ (light grey). Events resulting from TALEN-mediated DSB followed by HDR in the presence of the *yfp*\* repair template are shown in yellow. Cells expressing *gfp* only are not indicated. Each experiment was conducted on two independent transgenic plant lines. Bars represent the mean of five leaves (see Table 1).

In the present study, the first example is provided for plants, where a functional allele gains modified functionality after repair of TALEN-mediated DSBs. Townsend *et al*. (2009) have previously demonstrated by the ZFN approach using tobacco protoplasts that the introduction of point mutations in *SurA* or *SurB* causes a loss of binding capacity between the concerned gene products and the herbicides chlorsulfurone and imazaquin, which is associated with an advantage under selective conditions. More recently, the same approach was conducted using TALENs instead of ZFNs which resulted in an efficiency of 4% of allele replacement (Zhang *et al*. 2013). In maize it was shown that simultaneous expression of ZFNs and delivery of a simple heterologous donor molecule leads to precise targeted addition of an herbicide-tolerance gene at the intended locus in a significant number of isolated events (Shukla *et al*. 2009). Other previous examples of functional restoration of target genes rested on RGEN-mediated DSB induction without use of repair template (Feng *et al*. 2013; Jiang *et al*. 2013; Baltes *et al*. 2014) or on endonuclease I-*Sce*I the applicability of which is largely confined to its native DNA target sequence (Fauser *et al*. 2014).

Since somatic cells are diploid and therefore carry two copies of each gene, loss of GFP activity would require the loss-of-function of both copies. Therefore, mutation efficiency is easily under-estimated. A number of the *yfp*-expressing cells co-expressed *gfp*, but others clearly showed solely YFP activity (Figure 2B, C, left cell). To confirm that the genomic *gfp* sequence had been converted into the *yfp*, PCR was conducted using primers (Table S1) specific for *yfp* and the 35S promoter driving the selectable marker gene (*HPT*), the latter expression unit being present within the stably inserted T-DNA that also carries the *gfp* gene. Thus, an amplicon could only be generated if a TALEN-induced DSB had been processed by HDR, based on the *yfp* fragment supplied in the form of the repair template (Figure 3A). Sequencing of the fragments recovered from the amplicons unambiguously confirmed that the desired conversion had occurred in the genomic DNA (Figure 3B).

**Figure 3.**
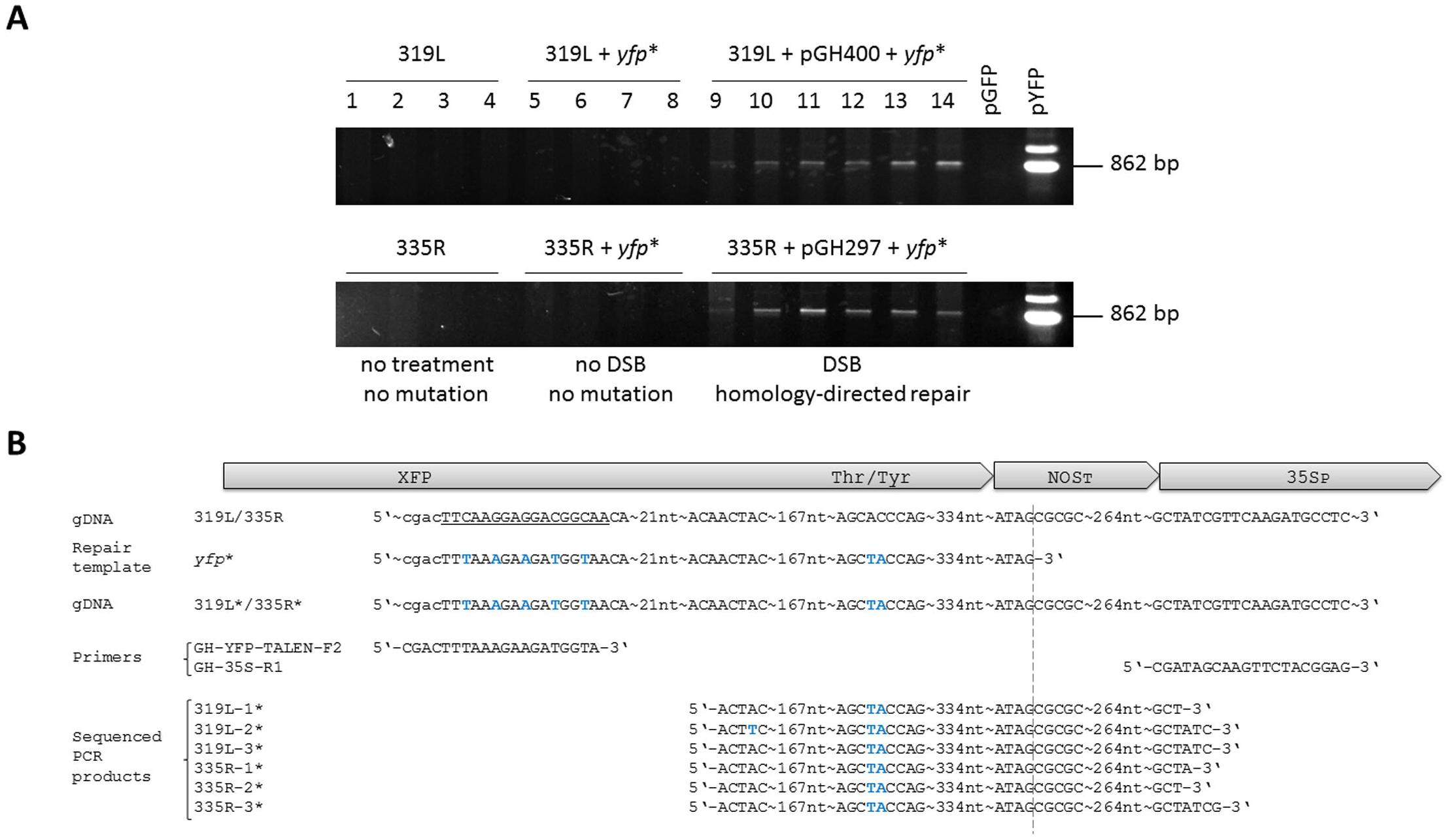
PCR analysis and DNA-sequencing confirm *gfp*-to-*yfp* conversion within the host genome. (A) PCR-products amplified from genomic DNA extracted from leaves of lines 319L (upper gel) and 335R (lower gel) without bombardment are shown in lanes 1 to 4, of those bombarded with the linearized truncated *yfp*\* fragment alone in lanes 5 to 8, and of those bombarded with both the linearized truncated *yfp*\* fragment and the complementary TALEN unit (pGH400 for right-hand, pGH297 for the left-hand unit) in lanes 9 to 14. pGFP was the negative control (plasmid carrying *gfp* T-DNA) and pYFP positive control (plasmid carrying *yfp* T-DNA). (B) Verification of allele conversion via sequencing; presented are sequences of donor plant (gDNA), the truncated *yfp*\* fragment used as repair template, the genomic target sequence after homology-directed repair of TALEN-mediated DSBs, forward and reverse primers used for DNA-amplification from samples of different leaves of lines 319L (left-hand TALEN unit) and 335R (right-hand TALEN unit) 20 h after bombardment as well as sequencing data providing evidence of *gfp*-to-*yfp* conversion. The sequences recovered from all other independent leaves were identical to those three per plant line shown here. Nucleotides different in the *yfp*\* sequence from those of *gfp* are printed in blue. The binding site of the right TALEN unit is underlined. An asterisk indicates an altered target sequence after leaf bombardment in lines 339L and 335R. The dotted line indicates the end of the repair template. XFP – green or yellow fluorescent protein, Thr/Tyr – amino acid change responsible for GFP-to-YFP conversion, NOST: A. tumefaciens NOPALINE SYNTHASE transcriptional termination sequence, 35SP: CaMV 35S promoter.

A challenge which remains to be overcome before the principle demonstrated here at the cellular level can be extended to the regeneration of plants carrying a precisely predicted modification is that such genome editing events are not typically associated with any selective advantage. As a result, alternatives to the conventional use of selective agents (such as herbicide or antibiotic resistance) will need to be elaborated. However, for the moment, what has been demonstrated is the proof of concept that any plant gene can be modified in a predictable way, thereby opening the door to redesigning native gene sequences to modify their function or to reverse mutations in a directed way.

Predictable genetic modifications comprising only a few genomic base pairs rather than entire genes are of particular practical relevance, because they might not fall under the European regulation of genetically engineered organisms. One prominent example where the precise editing of only a small genomic motif may result in a valuable trait is the wheat Lr34 ABC transporter which confers durable resistance to multiple fungal pathogens (Krattinger *et al*. 2009). The authors showed that resistance was associated with either the deletion of a phenylalanine residue (M19) or a single-nucleotide polymorphism in exon 12 which causes a tyrosine to a histidine (M21) conversion. Likewise, small alterations in the nucleotide sequence associated with valuable gain or modification of function have been shown in many other plant genes, e.g. those involved in resistance or flowering time (Rubens *et al.* 2006, Dally *et al*. 2014). The capacity to achieve this would represent a quantum improvement over established methods of random mutagenesis or the knock-in of an entire coding sequence. The technique promises to revolutionize the way plant genes are functionally validated, not to mention to create opportunities for crop improvement based on the paradigm that any sequence-defined allelic variant can be established at its native locus without perturbing the genetic background.

### Conclusion

TALEN-mediated targeted mutation has been previously shown in a variety of plant species. The results presented here provide evidence that targeted allele conversion predictably gaining a new function is also feasible. While previous reports showing that customized repair templates can be used for HDR of DSBs in plants have been largely confined to dicots like Arabidopsis or tobacco, a first example of this approach is given here using a Triticeae tribe member with high agronomic importance. In addition, a readily screenable marker system is introduced that might be useful for methodological optimization studies in the field of genome editing.

## Abbreviations

HDR: Homology-directed repair;
NHEJ: Non-homologous end joining;
RGEN: RNA guided genome editing nucleases;
GT: Gene targeting;
DSBs: Double strand breaks.

## Acknowledgements

We appreciate the excellent technical assistance of Petra Hoffmeister, Andrea Mueller and Ingrid Otto.

